# Beyond Microbial Abundance: Metadata Integration Enhances Disease Prediction in Human Microbiome Studies

**DOI:** 10.1101/2025.09.03.674104

**Authors:** Andre R Goncalves, Hiranmayi Ranganathan, Camilo Valdes, Haonan Zhu, Boya Zhang, Car Reen Kok, Jose Manuel Martí, Nisha J Mulakken, James B Thissen, Crystal Jaing, Nicholas A Be

## Abstract

Multiple studies have highlighted the human microbiome’s potential as a biomarker for diagnosing diseases through its interaction with systems like the gut, immune, liver, and skin via key axes. Advances in sequencing technologies and highperformance computing have enabled the analysis of large-scale metagenomic data, facilitating the use of machine learning to predict disease likelihood from microbiome profiles. However, challenges such as compositionality, high dimensionality, sparsity, and limited sample sizes have hindered the development of actionable models. One strategy to improve these models is by incorporating key metadata from both the host and sample collection/processing protocols. In this paper, we introduce a machine learning-based pipeline for predicting human disease states by integrating host and protocol metadata with microbiome abundance profiles from 68 different studies, processed through a common pipeline. Our findings indicate that metadata can enhance machine learning predictions, particularly at higher taxonomic ranks like Kingdom and Phylum, though this effect diminishes at lower ranks. Our study leverages a large collection of microbiome datasets comprising of 11,208 samples, therefore enhancing the robustness and statistical confidence of our findings. This work is a critical step toward utilizing microbiome and metadata for predicting diseases such as gastrointestinal infections, diabetes, cancer, and neurological disorders.

## 1 Introduction

Imbalances in the human microbiome have been associated with conditions such as inflammatory bowel disease (1), obesity (2), diabetes (3), cancer (4), and neurological disorders (5, 6). Distinct microbial profiles can influence disease progression and treatment responses, underscoring the microbiome’s potential as both a source of biomarkers and a therapeutic target. This potential paves the way for training machine learning (ML) models—given adequate training data—that use microbiome abundance information to predict the likelihood of a host developing specific diseases. Recent studies (7–9) have demonstrated the potential of ML techniques in this field. However, the development of actionable models can be hindered by challenges pertaining to microbiome data including (10): (1) limited sample sizes in individual studies; (2) the additional complexity arising from the compositional nature of microbiome data; (3) the high dimensionality and sparsity of microbiome abundance profiles, particularly at lower taxonomic levels such as Genus and Species; (4) biases introduced by variations in sample collection, processing, and sequencing protocols across studies; and (5) the absence of standardized datasets and protocols for evaluating ML models.

One strategy to improve machine learning models is by integrating key metadata from both host and sample collection/processing protocols. Host demographic factors, such as age and biological sex (11, 12), significantly influence microbiome composition, potentially altering disease susceptibility. Lifestyle and diet are crucial determinants of a healthy microbiome, while genetic predispositions can affect conditions like obesity, which, in turn, impact the gut microbiota (13). Additionally, details about sample origin, collection, and sequencing methodologies can help reduce dataset biases. However, a major challenge lies in the lack of standardized host metadata variables across studies, as these are often conducted independently with distinct research questions in mind. To develop a machine learning model that effectively incorporates host and sample metadata, a consistent set of variables must be available across all studies. In cases of missing data, imputation techniques can be applied to fill these gaps.

More effective incorporation of metadata variables in future efforts will guard against artifactually inaccurate conclusions and limit the impact of systematic confounding factors; indeed, previous reviews have specifically stressed the challenge and criticality of more effective, integrated metadata processing methods (14). Numerous important and laudable efforts are underway to improve metadata recording and curation, which are improving the accessibility of these datasets for model training (15–17). These efforts will be further strengthened with testing and adoption of validated methods for metadata imputation, which will also facilitate the use and integration of past and current studies with incomplete or sparse metadata provision.

In this paper, we introduce a machine learning-based pipeline for predicting human disease states by integrating host and protocol metadata with microbiome abundance profiles from 68 different studies. Because not all studies collected the same set of metadata variables, we employ data imputation strategies to effectively utilize data from all sources. We quantitatively assess the impact of metadata on model classification accuracy and discuss factors that can both increase and decrease model performance across different disease groups when the input feature set is expanded with metadata. Additionally, we analyze which metadata variables provide the most informative insights. To our knowledge, this study leverages the largest meta-study collection of microbiome datasets derived from shotgun metagenomic sequence data, with curated metadata features, available in the literature, thereby enhancing the robustness and statistical confidence of our findings.

## 2 Methods and Materials

### A Human Disease Prediction from Host Metadata and Microbiome Profile

Host disease prediction from microbiome profiles involves the application of machine learning models to classify individuals as either “diseased” or “control” based on their microbiome composition. These profiles are collected from specific body sites, depending on the interactions and mechanisms of interest between disease states and host microbiome sites. For example, skin or gut microbiome samples could be analyzed for dermatological conditions to either understand the direct effect of skin microbial communities on the disease or to study the bidirectional relationship of the gut-skin axis. The input to the machine learning models consists of normalized microbiome abundance profiles, which represent the relative abundances of microbial taxa in each individual. These profiles are preprocessed using normalization techniques to account for the compositional nature of microbiome data and mitigate the effects of varying sequencing depths. The centered log-ratio (CLR) transformation (18) is one of the most widely used methods and is applied in this study as well. The normalized profiles are then fed into the models, which are trained to predict whether a given microbiome profile is associated with a *diseased* or *control* individual, based on a specific disease under investigation. By identifying subtle microbial signatures, these models have the potential to detect disease early and provide valuable insights into underlying disease mechanisms. Figure 1 illustrates the pipeline employed in this study. Each step of the pipeline is explained in detail in the following sections.

**Fig. 1.**
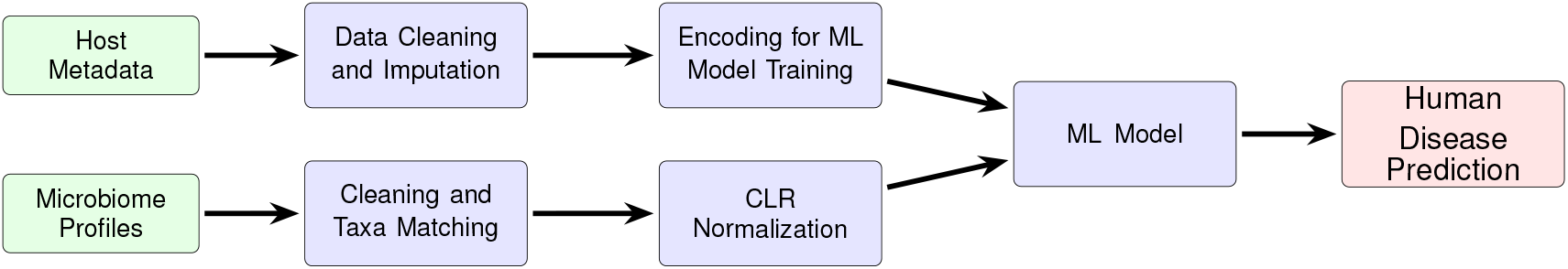
Pipeline for predicting human diseases using host metadata and microbiome profiles.

For this analysis, we evaluated four well-established machine learning models: k-Nearest Neighbors (KNN), Logistic Classifier (LC), Linear Support Vector Classifier (LinearSVC), and Random Forest (RF). While other non-linear methods were considered, their performance was suboptimal compared to these selected models, so they were not included in the final evaluation. All models were implemented using the Scikit-learn library (19).

### B Data

#### B.1 Data Collection and Processing

To facilitate microbiome machine learning, we leveraged high-performance computing (HPC) resources to perform metagenomic classification on publicly available metagenomic sequence data, in addition to robust metadata curation. This resulted in structurally consistent microbiome taxonomy feature count tables for 13,897 samples across 84 studies.

Methods of this pipeline were previously described (20, 21). Briefly, literature studies were selected according to defined intake criteria and each metagenomic dataset assigned a label of “control” or “diseased”. Metadata were extracted and a controlled vocabulary applied for variables such as disease state, adapting and, where possible, adhering to conventions established by the Genomic Standards Consortium (22). Raw sequence data (fastq files) were downloaded from the appropriate public repository, pre-processed via fastp (23) and defined threshold criteria. These metagenomic samples were then processed for taxonomic classification via Centrifuge (24) using a novel decontaminated index covering the entire tree of life (20) constructed from the NCBI BLAST Nucleotide (nt) database (25). Finally, taxonomic results were post-processed using Recentrifuge (26) to normalize classification scores and perform additional filtering.

From the library of 84 studies, 10 contained only samples with single-class labels “control” or “diseased”. Since our goal was to train classifiers on individual studies using supervised machine learning models, which require samples from both classes, these studies were excluded from our analysis. Additionally, some studies had very limited sample sizes or were extremely class imbalanced. In both cases, it is challenging to accurately assess classification performance. Therefore, we also excluded studies with fewer than 20 samples or with a class imbalance greater than 98%. Ultimately, we retained a set of 68 microbiome studies, as shown in Figure 2, along with basic statistics for each study. The full list of referenced microbiome studies can be found in the Supplementary Material.

**Fig. 2.**
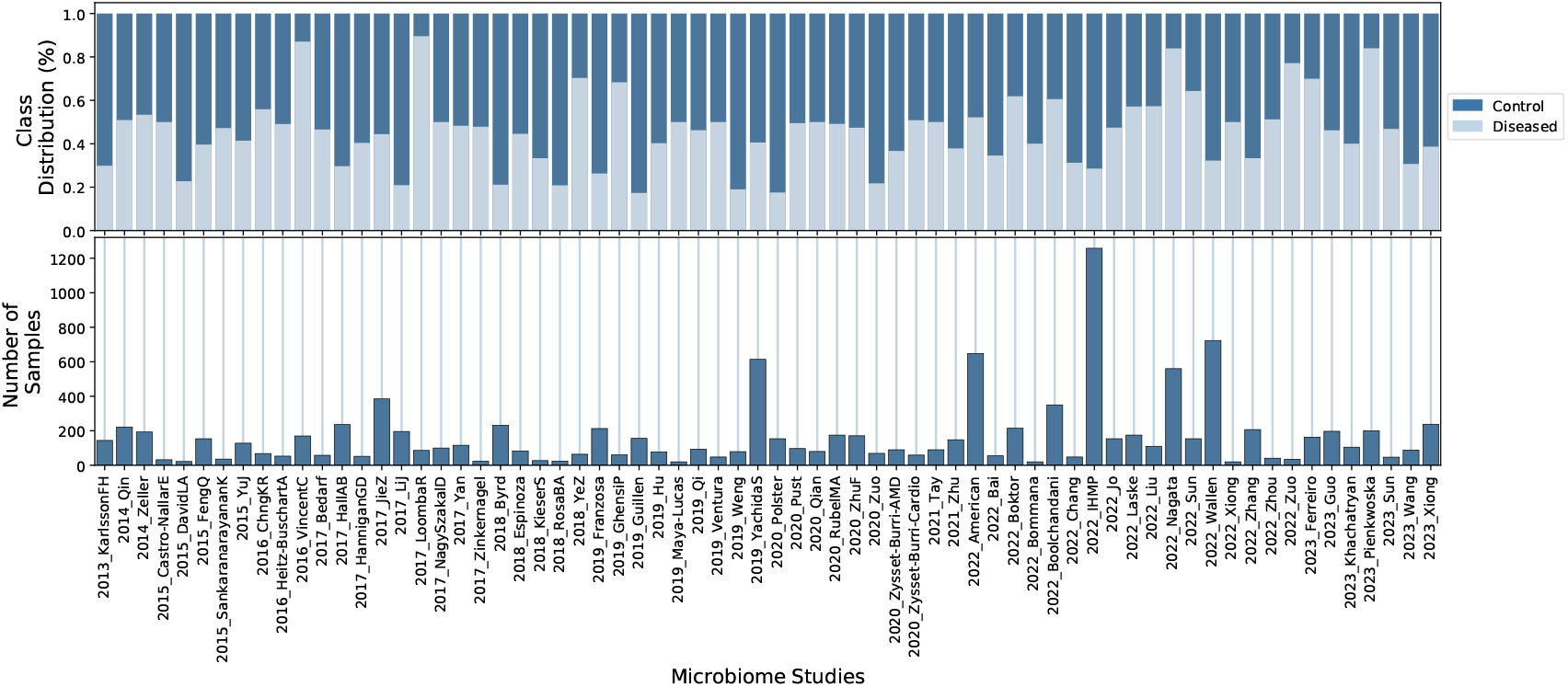
Summary statistics for the 68 microbiome studies used in this work. The bottom plot displays the number of samples in each study, while the top plot shows the proportion of control versus diseased individuals in each study. Additional details and references for the microbiome studies can be found in the Supplementary Material.

**Fig. 3.**
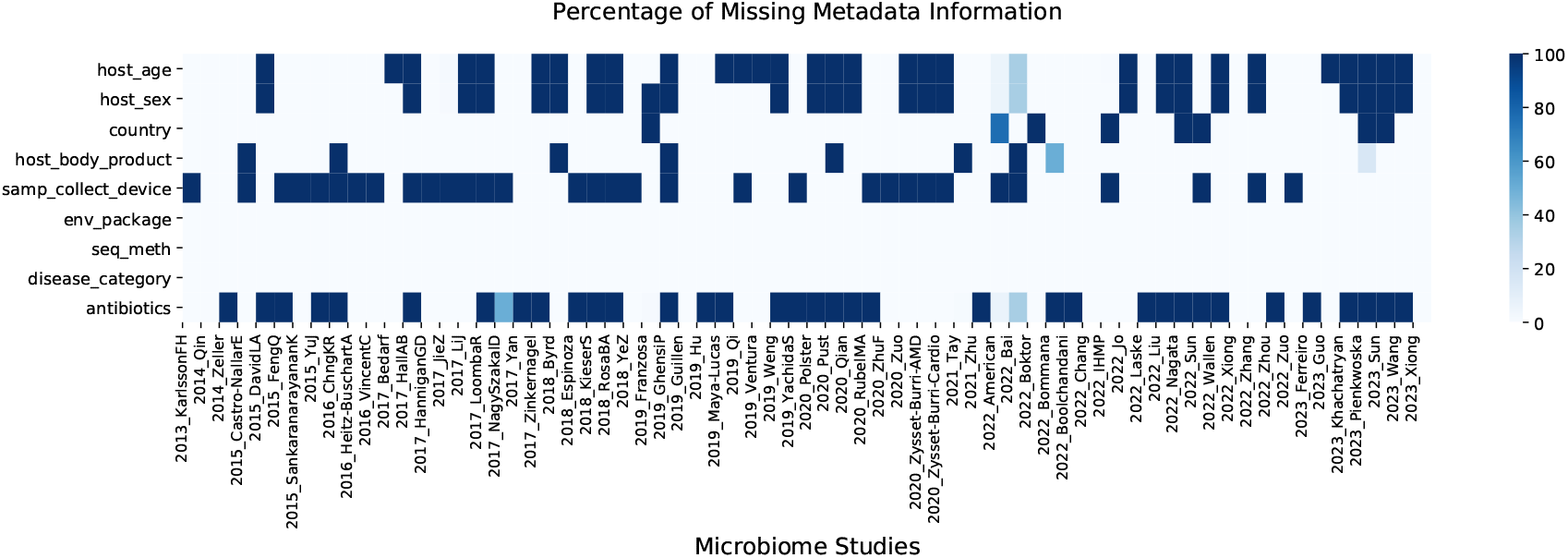
Percentage of missing metadata information for each variable across all studies.

#### B.2 Metadata Variable Selection

The union of metadata variables from the 68 microbiome studies comprised 171 variables. As expected, not all variables were collected in all studies, resulting in a large portion of the 171 variables being absent in many studies. Missing data significantly influences which subset of metadata is ultimately retained for use in machine learning models, as it can drastically affect the performance of the methods, a topic we will discuss in Section B.3.

We employed a variable selection strategy that considers both the percentage of missing data for each variable in all studies and the variable’s predictive power. There is a clear trade-off when retaining variables with missing data: while they may contain valuable subject information, they can also introduce noise if data imputation is performed.

From the original set of 171 variables, we selected a total of 9 predictive variables listed in Table 1. The selection process is described below. For the `disease_category` variable, we consolidated related diseases into broader categories to streamline the model, effectively reducing the number of disease categories without compromising predictive precision. The steps in metadata feature selection and the number of remaining features after each step are shown as a flow chart in Fig. 4.

**Table 1.** Characteristics of the retained host metadata variables. ^*^The number of categories is aggregated across all studies, including the ‘UNK’ category.

| Name | Description | Type | # of Categories* |
| --- | --- | --- | --- |
| host_age | Host age. | Numerical | - |
| host_sex | Host biological sex. | Categorical | 3 |
| country | Geographical location. | Categorical | 33 |
| host_body_product | Sampled body location. | Categorical | 7 |
| samp_collect_device | Device used to collect sample. | Categorical | 9 |
| env_package | Sample source description. | Categorical | 4 |
| seq_meth | Sequencing method used. | Categorical | 12 |
| disease_category | Type of disease. | Categorical | 22 |
| antibiotics | Antibiotics use. | Categorical | 3 |

**Fig. 4.**
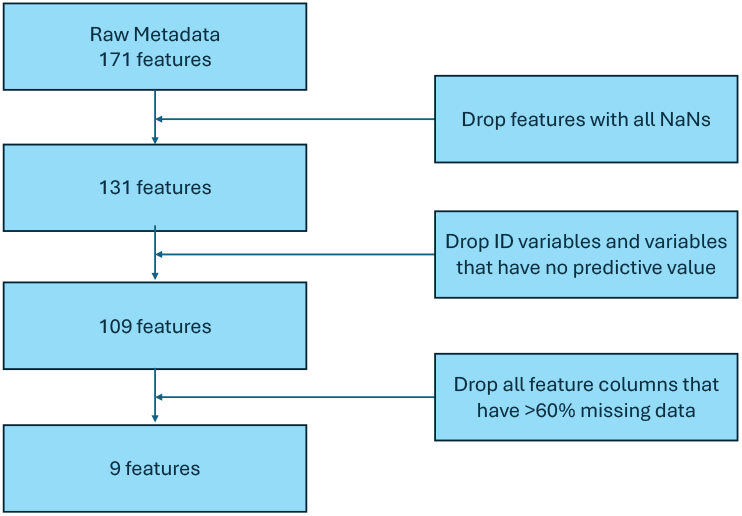
Flowchart for metadata feature processing.

#### B.3 Missing Metadata Imputation

For host metadata variables to be used in machine learning models, missing values (NAs/NaNs) must be assigned a numerical representation. Various strategies have been developed in the field of data imputation. In this study, we employ and compare three different imputation techniques:

1. Imputation of all missing variables using the MICE algorithm (27) (`MICE_ALL`);
2. Imputation of `host_age` using MICE with UNK (unknown) for missing data in other categorical variables (`UNK_MICE_Host_Age`);
3. Imputation of `host_age` using k-NN on the microbiome profiles with UNK for missing data in other categorical variables (`UNK_KNN_Host_Age`).

The retained metadata variables include a combination of subject-specific (e.g., age, sex) and protocol-specific (e.g., body location, sequencing method) variables. While subject-specific variables are most relevant to indicating and discriminating health state, variables describing the collection and analytical methods may be influential with respect to systematic distinctions in observed microbial features between studies. As such, they may be instructive in this and future studies, and are included in the describe assessment. Since host age is a key factor influencing microbiome composition (28–30), it is expected to play a significant role in the machine learning model. Therefore, we place special emphasis on accurately imputing this variable.

In the first imputation method, we use MICE (27, 31). MICE stands for Multivariate Imputation by Chained Equations. It is a statistical method used to handle missing data in a dataset. The process uses multiple imputation techniques to fill in the missing data and then combines the results from multiple imputations to produce a final imputed dataset. We impute all variables using MICE and call this `MICE_ALL`. As a second method of imputation, we impute `host_age` using MICE and mark missing data in other categorical variables as UNK. We call this method `UNK_MICE_Host_Age`. For the third method, we impute `host_age` using k-NN on the microbiome profiles and label missing data in other categorical variables as UNK, a method we refer to as `UNK_KNN_Host_Age`. Here, a new sample is imputed by identifying the closest samples in the training set and averaging their values to fill in the missing data. In our experiments, we used *k* = 1, but we also evaluated *k* = 3 and *k* = 5 and observed no significant differences.

#### B.4 Feature Encoding

Machine learning algorithms require that all predictive variables be represented numerically. This is called *feature encoding*. We employ different encoding strategies for different types of predictive variables. The selection depends on the characteristics of the variables under consideration. In what follows, we use the terms variable and feature interchangeably.

Numerical variables are already numbers and can be used directly as input features to machine learning models. Therefore, `host_age` does not need any additional encoding. Variables that are purely categorical (qualitative variables with no clear ordering or associated numerical values) are usually represented via one-hot encoding. In this encoding strategy, the variable is represented by a binary vector of the same size as the number of categories. The vector contains zeros everywhere except for the position corresponding to the given category, in which case a 1 appears. This is equivalent to introducing dummy variables for categorical variables in standard regression analysis. All categorical variables in Table 1 were encoded using this strategy.

Cases with missing categories in `samp_collect_device`, `disease_category`, `host_sex`, `antibiotics`, `host_body_product` and `country` are treated as an additional category for that feature, unless specified otherwise. We ignore all features that are not directly related to host condition such as IDs, study abstracts, paper references, etc.

#### B.5 Outcome Definition

For each individual study, a binary variable indicating “control” (0) or “diseased” (1) serves as the outcome variable for the corresponding machine learning model. It is important to note that the definition of “control” is specific to each study. For instance, in a gastrointestinal (GI) study, a “control” label means the individual did not exhibit any GI disorders at the time of sample collection; however, other conditions like diabetes, cancer, or neurological disorders could theoretically be present but were not assessed. Similarly, a sample classified as “diseased” in a GI study may be considered “control” in the context of a pulmonary disease study.

This suggests that disease state classification is better approached as a multi-label problem, since an individual may have multiple coexisting conditions. However, the existing collection of microbiome datasets does not support this approach as the underlying studies assessed a single given disease condition. Furthermore, the variability and specificity of disease definitions across studies make it difficult to train a single, generalizable model to differentiate between control and diseased states using data from all studies combined, even though this would increase the training sample size. Therefore, developing multi-label models or broadly generalizable models is beyond the scope of this article.

#### C Experimental setup

For each ML model in each microbiome study, we conducted 30 independent experiments using holdout cross-validation with varying train/validation/test splits. We allocated 70% of the data for training, 10% for validation, and 20% for testing. The hyperparameters used for each model are detailed in the Supplementary Material. The host disease prediction performance is evaluated using the widely recognized area under the Receiver Operating Characteristic curve (AUROC) metric (32). This metric assesses the trade-off between the True Positive Rate (Sensitivity) and the False Positive Rate (1 - Specificity) across various threshold settings, providing a comprehensive measure of the model’s discriminatory ability.

## 3 Results and Discussion

### A Host Metadata Only

We first investigated and compared the metadata imputation models presented in Section B.3. For this, we trained binary classification models to distinguish between diseased and control patients using *only metadata*. The goal of this analysis was to assess the impact of each imputation method, as the microbiome profiles will be incorporated into the feature vectors in subsequent steps. Figure 5 presents the classification performance using host metadata as input. While the overall performance is modest, the models were able to extract some relevant discriminative information from the metadata (with Random Forest achieving over 62% accuracy) and the selected metadata variables alone are not necessarily expected to comprehensively indicate, on their own, disease state. Notably, the imputation technique based on the nearest neighbor in microbiome space led to better classification performance compared to the other two approaches. Given its improved performance, we will only consider the `UNK_KNN_Host_Age` imputation method for the remainder of this analysis.

**Fig. 5.**
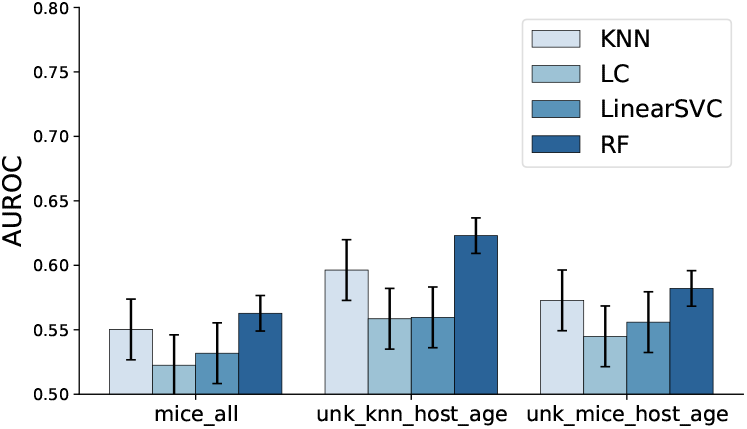
Human disease classification performance using *only host metadata* information as discussed in Sections B.2, B.3, and B.4.

### B Microbiome Only

Figure 6 presents the average performance across the 30 runs for each study. The error bars represent the variance in the ML model’s performance at each taxonomic level across all studies. As can be seen, the classification performance improves noticeably as we move to lower taxonomic levels (from Kingdom to Species). However, working with profiles at these finer taxonomic levels demands significantly more computational resources due to the increased dimensionality of the data (e.g., microbiome abundance profiles have 11 dimensions at the Kingdom level but nearly 31,000 at the Species level, as these profiles are derived from shotgun metagenomic data across all domains). The high dimensionality not only increases the time required for data processing and normalization but also leads to substantially longer training times for machine learning models. Overall, the models demonstrate better performance when microbial abundance is analyzed at lower taxonomic levels. This is consistent with previous observations that classification performance generally improves with increasing taxonomic resolution (33).

**Fig. 6.**
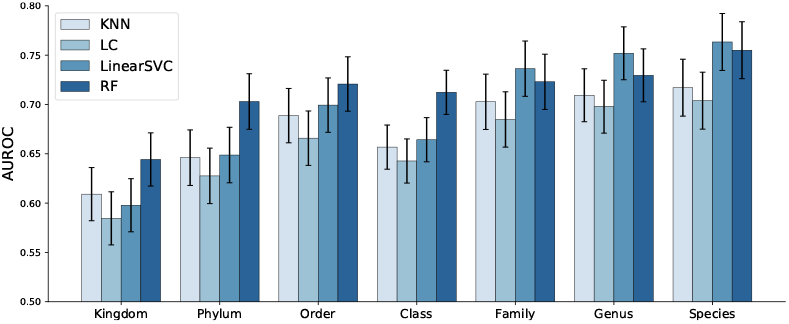
Classification performance (AUROC) of the ML models across different taxonomic levels, based *solely on microbiome profile* information. Overall, the models demonstrate better performance when microbial abundance is analyzed at lower taxonomic levels.

In Figure 6, we also highlight the performance of the ML methods, showing that the LinearSVC and RF are the top performing models. RF slightly outperformed LinearSVC at higher taxonomic levels (Kingdom, Phylum, Order, and Class), while LinearSVC yielded more accurate predictions at lower taxonomic levels (Family, Genus, and Species). This could be a reflection of the relative flexibility of RF to operate across nonlinear decision boundaries resultant from relationships between coarser taxonomic ranks. This is consistent with previous model comparisons for inflammatory bowel disease (34). It may be that, in this case, the LinearSVC more effectively processes the sparser, higher-dimensional species level features.

### C Host Metadata and Microbiome Jointly

Classification performance of ML models using both host metadata and microbiome profiles as input is presented in Figure 7. Notably, the Random Forest model demonstrates strong performance even at higher taxonomic levels, such as Kingdom and Phylum, highlighting the importance of incorporating host metadata. As can be seen in Figure 8 the improvement affiliated with metadata inclusion decreases as we move to lower taxonomic levels and becomes negligible, in the current analysis, at the Species level. A likely explanation for this diminishing improvement is that the influence of metadata features diminishes as the problem’s dimensionality increases from dozens to tens of thousands. In other words, the limited metadata variables become “diluted” by the extensive microbiome metagenomic profile information.

**Fig. 7.**
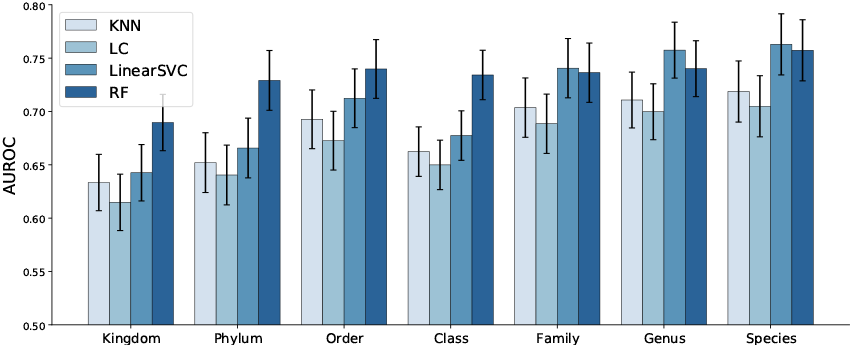
AUROC performance of the ML models across various taxonomic levels, utilizing both *host metadata and microbiome profiles*.

**Fig. 8.**
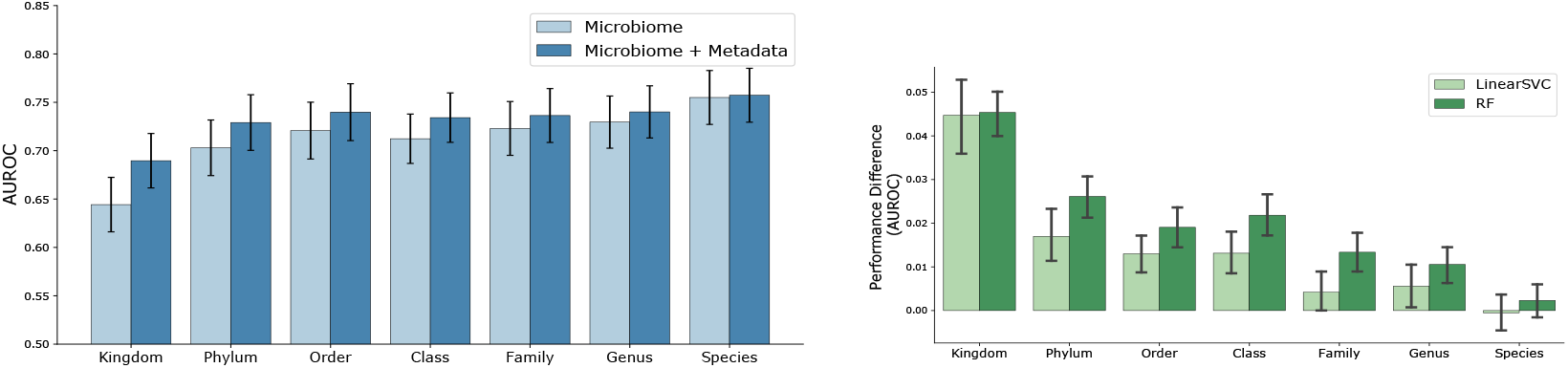
Comparative classification of ML models trained using *microbiome only* vs *microbiome + host metadata*. Left figure shows the AUROC for RF, and right figure presents the difference in performance for both RF and LinearSVC model. Significant improvement can be observed in particular for higher taxonomic levels.

It is also important to consider the impact of the type of machine learning model used, as different models handle features in distinct ways. Some models, like k-NN, logistic regression, and LinearSVC, evaluate all features simultaneously, often by considering the distances between entire feature sets. In contrast, others, such as Random Forest, can focus on specific subsets of features, particularly in tree-based models with limited tree depth. This distinction can significantly influence model performance and interpretation when a limited set of metadata is incorporated into complex and high-dimensional microbiome profile data. In that sense, models with feature-level focus can more readily pick up the limited metadata features. Figures 8 show a comparative performance plot for both Random Forest and LinearSVC, respectively. For the Random Forest model, the improvement of microbiome plus metadata over microbiome only is more noticeable than LinearSVC’s for the majority of the taxonomy levels, in particular for lower levels such as Family and Genus.

### D Analysis at Disease Group Level

Given the varying degrees of association between microbiome profiles and specific diseases, as well as the differing significance of host metadata across disease categories, it is reasonable to hypothesize that incorporating host metadata may significantly enhance classification performance for certain disease groups more than others. This analysis is essential to identify where host factors play a pivotal role alongside the microbiome in disease prediction, as some diseases may be more influenced by host characteristics such as age, sex, or antibiotic usage, for instance, distinctions in immune response mediated by gender (12).

In Figure 9, we present the classification performance difference between the *host metadata + microbiome* and *microbiome only* RF models for each study for the Kingdom, Order and Species taxonomic ranks. The plots depict the mean and 95% confidence interval of the performance improvements resulting from the addition of metadata variables to the microbiome metagenomic profiles, across independent runs. This detailed comparison allows us to assess the specific impact of host metadata on the predictive accuracy of the models across various disease groups, providing insights into the contexts where host information is most beneficial.

**Fig. 9.**
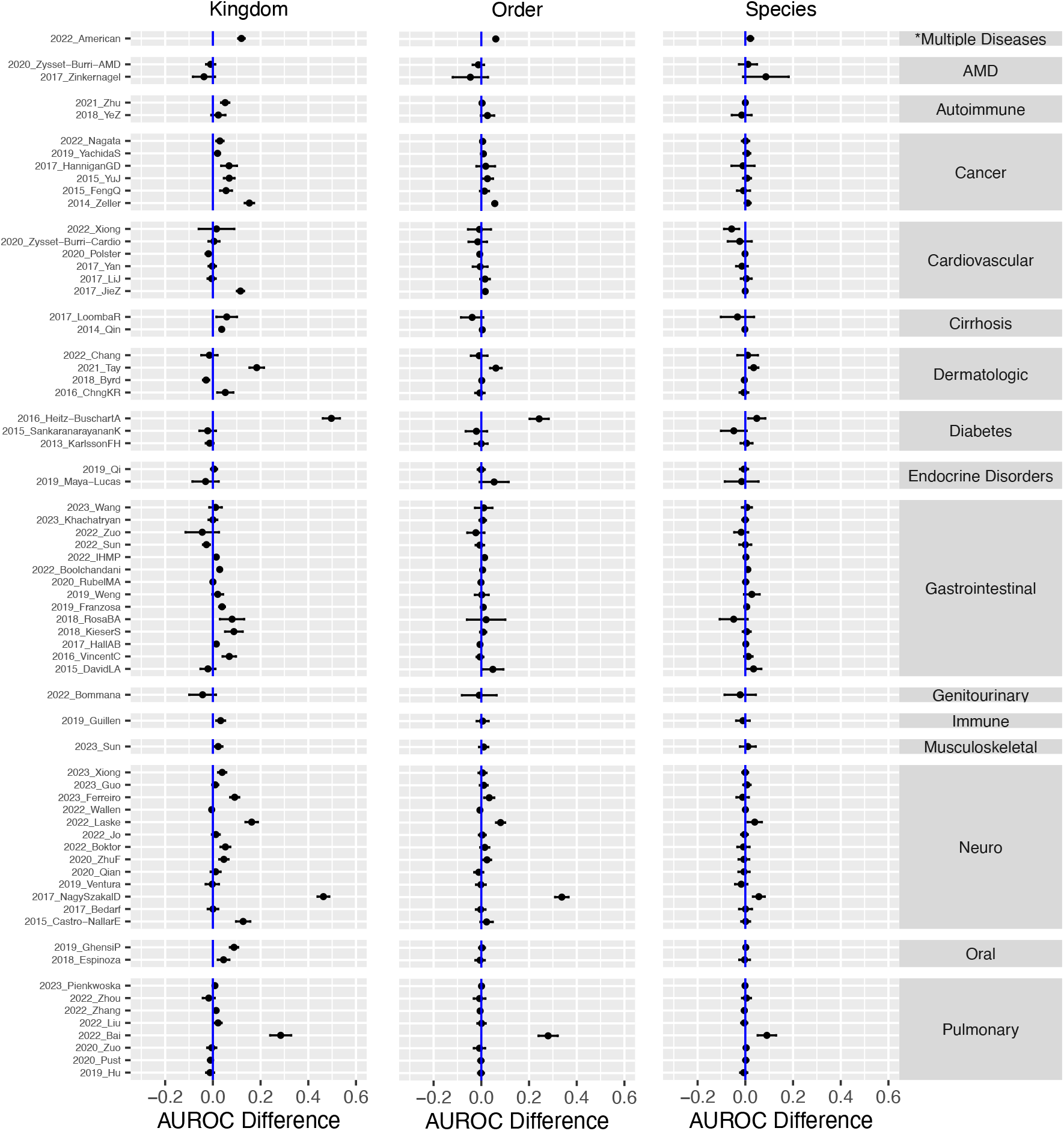
Performance differences in human disease classification between *microbiome + host metadata* and *microbiome only* across all studies, grouped by disease category. Positive values indicate improved performance with the inclusion of host metadata. The plot shows the mean and 95% confidence interval of the RF’s AUROC difference. *The American Gut Project (`2022_American`) was a crowdsourced study with diverse participants having various health conditions rather than focusing on one specific disease.

As previously observed in Figures 8, host metadata exerts a more pronounced effect at higher taxonomic ranks, such as Kingdom and Phylum, compared to lower ranks like Genus and Species. We hypothesize that this phenomenon is due to the “dilution” of metadata effects within the more complex, higher-dimensional, and potentially more physiologically representative microbiome abundance features at more specific taxonomic resolution.

At the Kingdom level, Figure 9 shows that host metadata significantly enhances classification performance for the Neuro (13 studies) and Cancer (6 studies) categories. In these groups, incorporating host metadata consistently improves or at least maintains performance across all studies, highlighting a strong and robust effect. This is likely attributable to specific metadata variables that are known to be associated with epidemiology of these diseases. Cancer-related studies included primarily assessments of colorectal cancer, with one study of pancreatic cancer. Age (35), gender (36), and antibiotic usage (37) are all variables influencing occurrence and risk for colorectal cancer. The neurological studies employed here included evaluations of Alzheimer’s disease (AD), Parkinson’s disease (PD), neuropsychiatric disorders, myalgic encephalomyelitis, multiple sclerosis. Similarly, such metadata variables are affiliated with the neurological diseases described, particularly for AD (38).

Other disease categories, such as Autoimmune, Cirrhosis, and Oral, also show noticeable performance gains, though the smaller number of studies in these categories may limit the generalizability of these observations. In contrast, larger disease groups like Cardiovascular (6 studies) and Pulmonary (8 studies) show virtually no performance change, with only one study in each group showing improvement. This suggests that additional or different types of host metadata might be needed for these disease groups, warranting further investigation into other host factors. It is also possible that the broad range of disease conditions categorized within the cardiovascular and pulmonary groups occludes the capacity of the current analysis to identify and employ consensus metadata variables with utility. For instance, studies specific to ischemic heart disease (39) would likely demonstrate clearer benefit.

For the largest group, Gastrointestinal (14 studies), the majority of studies exhibit improved performance with the inclusion of metadata. However, a few studies show no change, and some even display a decline in performance. It is important to note that this category encompasses a heterogeneous set of conditions, including diarrhea, gastrointestinal infections, irritable bowel syndrome, inflammatory bowel disease, and seasickness, which, as noted above, likely contributes to the variable impact of metadata inclusion.

### E Missing Host Metadata and Disease Prediction Performance

Figure 9 prompts a critical question: what factors contribute to performance improvement in some studies while leading to a decline in others, and in which cases is data imputation measurably improving performance? Several factors could be at play, including the relevance of the specific set of host metadata variables available and the sample size. However, a key factor is the extent of missing host metadata, which was subsequently imputed using the methods described in Section B.3.

Although imputing missing data enabled the use of host metadata in the ML model and generally improved performance, it also introduced noise into the final feature set. This is an expected result, and imputation can potentially introduce bias and obscure true biological signals (40). Our imputation techniques leveraged machine learning models to minimize this noise and maximize the information derived from similar studies, consistent with observations that model-based methods reduce false positive results after imputation (41), yet some residual noise remains inevitable.

To quantitatively evaluate the impact of imputed missing data, we calculated the percentage of missing values for each host metadata variable across all studies and then aggregated these percentages for two groups: studies where host metadata had a positive impact and those where it had a negative impact. These aggregated percentages are presented in Table 2. The last column shows the percentage of missing data when all variables are considered.

**Table 2.** Average percentage of metadata information imputed over all studies grouped by studies in which metadata information had a Positive and Negative impact on human disease diagnosis (AUROC). Table shows percentage for each metadata variable with missing data and the mean percentage over all metadata variables.

|  | samp_collect_device |  | host_body_product |  | country |  | antibiotics |  | host_sex |  | host_age |  | Overall |  |
| --- | --- | --- | --- | --- | --- | --- | --- | --- | --- | --- | --- | --- | --- | --- |
|  | Positive | Negative | Positive | Negative | Positive | Negative | Positive | Negative | Positive | Negative | Positive | Negative | Positive | Negative |
| Kingdom | 51.0% | 40.7% | 11.9% | 9.2% | 13.2% | 9.1% | 50.2% | 53.6% | 33.4% | 46.2% | 36.8% | 56.0% | 32.8% | 35.8% |
| Phylum | 49.2% | 45.8% | 12.4% | 8.6% | 14.1% | 9.2% | 48.8% | 55.9% | 31.5% | 48.1% | 36.6% | 54.3% | 32.1% | 37.0% |
| Order | 48.3% | 46.6% | 12.2% | 9.0% | 13.8% | 9.5% | 50.0% | 53.6% | 34.0% | 42.3% | 38.2% | 49.4% | 32.7% | 35.1% |
| Class | 48.7% | 45.6% | 12.2% | 9.1% | 13.7% | 9.5% | 49.2% | 54.0% | 31.8% | 44.9% | 37.6% | 50.6% | 32.2% | 35.6% |
| Family | 50.3% | 43.3% | 11.4% | 10.0% | 12.5% | 10.5% | 48.5% | 55.2% | 32.6% | 44.4% | 37.7% | 50.9% | 32.2% | 35.7% |
| Genus | 47.6% | 47.2% | 12.5% | 9.0% | 13.5% | 9.6% | 49.5% | 54.5% | 35.1% | 40.9% | 39.4% | 48.2% | 32.9% | 34.9% |
| Species | 47.9% | 47.4% | 11.5% | 10.0% | 12.5% | 11.1% | 52.1% | 50.3% | 36.9% | 37.6% | 42.1% | 43.9% | 33.9% | 33.4% |

A key observation from Table 2 is that, when considering all host metadata variables (“Overall”), studies with a positive impact had lower levels of missing data across all taxonomic ranks, with a reduction of up to 5% at the Order level. This suggests that the more complete the host metadata, the greater the improvement in disease prediction performance, and limitations in the introduction of additional noise.

At the level of individual host metadata variables, an opposite trend is observed for `samp_collect_device`, `host_body_product`, and `country`, where studies with a positive impact exhibit a higher level of missing or imputed data. This can be explained by the RF model feature importance shown in Figure 10, which indicates that these three variables have little or no predictive power for host disease state. As a result, the model places minimal weight on them, regardless of the percentage of missing data. This observation is consistent with expectations, as these methodological parameters would be consistent across healthy and disease state samples within a given study. While it is clear that extraction method, library preparation, and sequencing platform can introduce biases in resultant microbiome profiles (42, 43), such biases are expected to be uniform across the labels implemented for classification in the current study. In contrast, variables such as `host_age`, `host_sex`, and `antibiotics` are highly relevant, as evidenced by their predictive value. Table 2 reveals that for these key variables, studies with a positive impact had significantly less missing data—up to 20% less compared to those with a negative impact. This strongly suggests that reduced performance in some studies can be attributed to the increased amount of missing data for these critical variables. It also underscores the importance of acquiring a complete set of metadata features, particularly those with high predictive value for the ML model. Variables affiliated with sample processing, including sequencing method, body location, and collection device, demonstrated minimal feature importance values. Again, this observation is anticipated as, for a given study, these variables should be consistent across control and disease specimens. However, the described framework may be useful in future assessments evaluating distinct methods in pooled specimen sets, where it may be desirable to integrate datasets from sample cohorts that underwent distinct processing workflows.

**Fig. 10.**
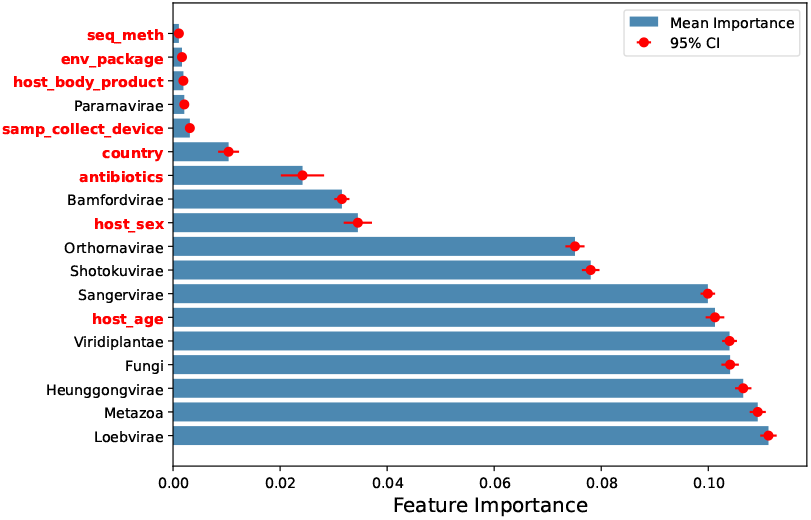
Random Forest model feature importance at Kingdom level. Host metadata variables are highlighted in red.

## 4 Conclusions

The broad range of consistency with which microbiome metadata are recorded and curated in the literature can render it difficult to perform consistent, ML-driven meta-studies that include these variables. In fact, without effective imputation strategies, the study presented here would not have been possible. We presented a pipeline that integrates microbiome abundance profile information and host and sample processing metadata to predict human disease state. These models can assess an individual’s potential resilience and provide the predicted probability of being “diseased”. Our multi-study investigation, comprising 68 published microbiome studies, reinforces the growing body of evidence supporting that incorporating metadata as additional inputs significantly improves the accuracy of disease prediction models. In our analyses this effect diminishes at lower taxonomic levels, such as Genus and Species. This may be attributed to the dilution of the limited metadata feature set compared to the extremely high dimensionality at these taxonomic levels, suggesting that additional metadata information could be required. Additionally, while missing metadata can be imputed using various methods, the noise introduced by imputation for critical variables can negatively impact performance for certain studies or disease groups. Impact on performance will also be dependent on the classifier method being applied.

Future efforts that should be prioritized for advancing ML-based microbiome analytics include in-depth investigation into the impact of metadata on model generalization to new, unseen studies. Such analyses would assess models trained on data from the same disease category as the new dataset, due to the varying definitions of “control” and “diseased” subjects across different studies, as discussed in Section B.5. Another potential avenue for exploration is the application of bias correction methods prior to training the ML models.

From our results, consistent with calls from numerous other consortia, we encourage the development of a standardized metadata collection framework for microbiome studies to ensure consistency and comparability across datasets. Such a framework would help address variability in data collection practices, enhance reproducibility, and ultimately support the development of more robust and generalizable machine learning models. The combination of uniform guidelines for metadata reporting with robust methods for data imputation would provide a framework that facilitates cross-study analysis and improves the model’s ability to generalize to new, unseen datasets, thereby advancing the capacity to generate predictive platforms based on microbiome-derived features.

## 6 Author Contributions

ARG: Conceptualization, Investigation, Methodology, Formal analysis, Software, Writing – original draft. HR: Conceptualization, Formal analysis, Writing – original draft. CV: Methodology, Writing – review & editing. HZ: Methodology, Writing – review & editing. BZ: Methodology, Writing – review & editing. CRK: Data curation, Writing – review & editing. JMM: Data curation, Writing – review & editing. JBT: Data curation, Writing – review & editing. CJ: Project administration, Funding acquisition. Writing – review & editing. NAB: Project administration, Funding acquisition, Writing – review & editing.

## 7 Acknowledgments and Funding

This work was performed under the auspices of the U.S. Department of Energy by Lawrence Livermore National Laboratory under Contract DE-AC52-07NA27344 and was supported by the LLNL Laboratory Directed Research and Development Program under Project No. 22-SI-002. Document release number LLNL-JRNL-2010890.

## 8 Data Availability Statement

The datasets utilized in this study can be found in https://gdo-meta2db.llnl.gov.

## Supplementary Material

### A Microbiome Studies

Table 3 include references to all the studies used in this work along with their metadata. Some studies were obtained from the *curatedMetagenomicData* package (44).

**Table 3.**
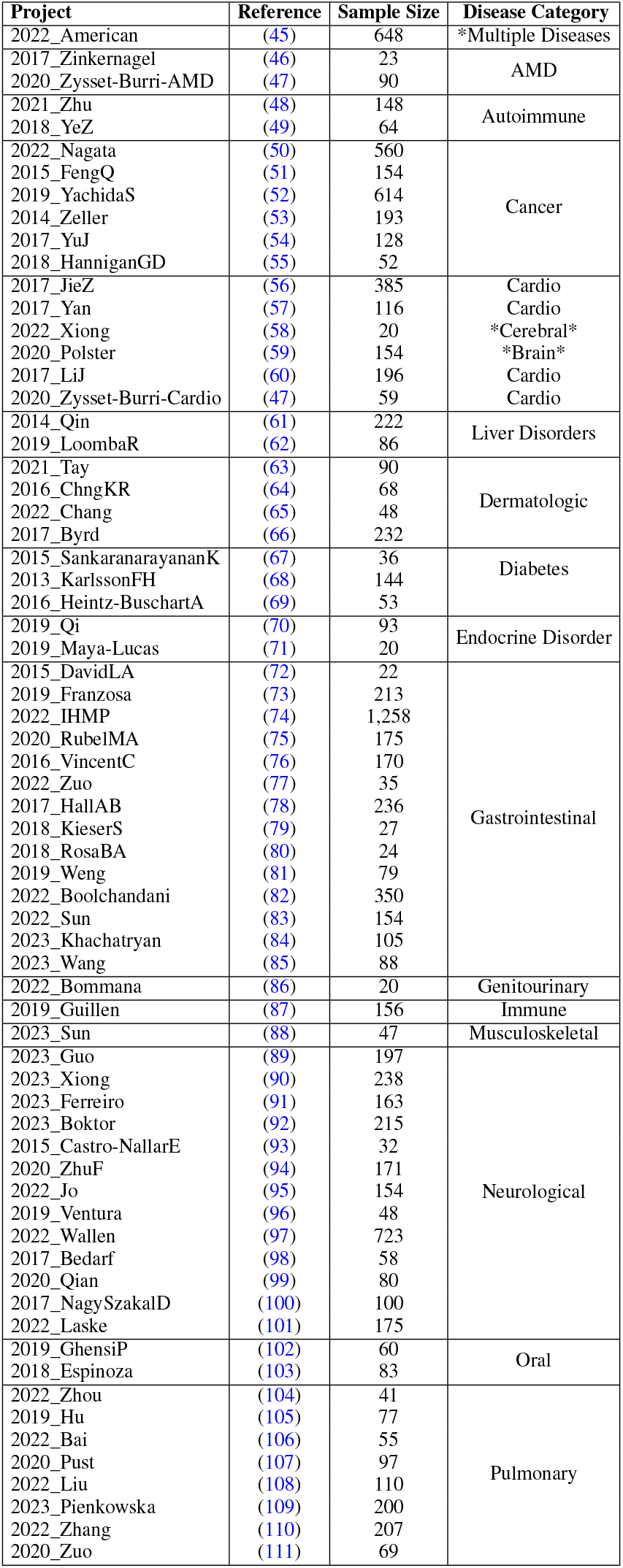
Microbiome studies used our meta-analysis work. We used a total of 11,208 microbiome profiles. *The American Gut Project was a crowdsourced study with diverse participants having various health conditions rather than focusing on one specific disease.

### B Machine Learning Model Hyperparameters

Table 4 summarizes the hyperparameter values applied for each machine learning algorithm in our study. These values were the same across all taxonomic ranks.

**Table 4.**
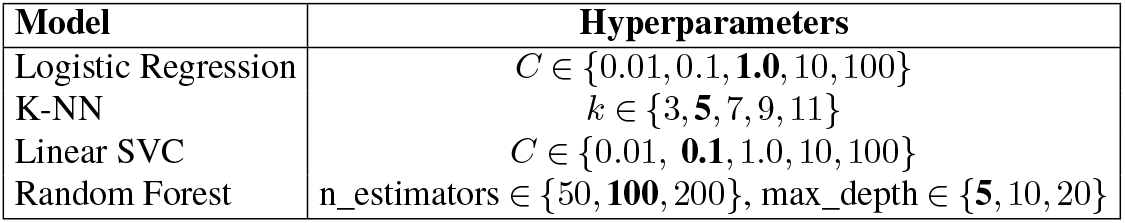
Hyperparameter values evaluated for each machine learning model via cross-validation. Optimal hyperparameters (highlighted in bold) were selected based on validation set performance and used for final test set evaluation.

